# Polymer-mediated oligonucleotide delivery enables construction of barcoded 3D cultures for spatial single-cell analysis

**DOI:** 10.1101/2023.11.20.567985

**Authors:** Jessica J. King, Alireza Mowla, Jessica A. Kretzmann, Marck Norret, Ulrich D. Kadolsky, Munir Iqbal, Alka Saxena, Somayra S.A. Mamsa, Sebastian E. Amos, Yu Suk Choi, Brendan F. Kennedy, K. Swaminathan Iyer, Nicole M. Smith, Cameron W. Evans

## Abstract

Spatial transcriptomics has been widely used to capture gene expression profiles, realised as a two-dimensional (2D) projection of RNA captured from tissue sections. Wree-dimensional (3D) cultures such as spheroids and organoids are highly promising alternatives to oversimplified and homogeneous 2D cell culture models, but existing spatial transcriptomic platforms do not currently have sufficient resolution for robust analysis of 3D cultures. We present a transfection-based method for fluorescent DNA barcoding of cell populations, and the subsequent construction of spheroidal cellular architectures using barcoded cells in a layer-by-layer approach. For the first time, changes in gene expression throughout this 3D culture architecture are interrogated using multiplex single-cell RNA sequencing in which DNA barcodes are used to encode the spatial positioning of cells. We show that transfection with fluorophore-conjugated barcode oligonucleotides enables both imaging and sequencing at single-cell resolution, providing spatial maps of gene expression and drug response. Additionally, we show that fluorophore-conjugated DNA barcodes support correlative imaging studies such as mechano-microscopy to capture information about spatially-varying mechanical heterogeneity in 3D cultures. We ability to create customised, encoded cellular assemblies is a general approach that can resolve spatial differences in gene expression in 3D cell culture models.

## Introduction

Single-cell RNA sequencing (scRNA-seq) has revolutionised transcriptomics for heterogeneous multicellular systems, enabling gene expression to be profiled within cell sub-populations. Cell hashing techniques for pooling multiple samples (‘multiplexing’) in one sequencing run has driven wider implementation of scRNA-seq, increasing throughput and reducing cost, manual labour and background noise^1^. To multiplex samples, techniques like CITE-seq^2^ and MULTI-seq^3^ couple unique barcode sequences with antibodies or lipids respectively, labelling cells with non-genomic DNA barcode sequences *via* interaction with cell surface antigens and the plasma membrane. In contrast, our recently reported single-cell transfection-enabled cell hashing and sequencing method, scTECH-seq^4^, uses a cationic polymer to deliver DNA barcodes into cells *via* endocytosis. Once barcoded, cells from different samples are pooled in one scRNA-seq run, during which the DNA barcodes are also sequenced, enabling cells to be demultiplexed and assigned to their originating sample(s). However, these existing barcoding methods do not contain spatial information *per se*, since cells are typically only labelled with barcode sequences after harvesting and cell dissociation, and prior to library preparation.

Engineered three-dimensional (3D) cultures in which information about cell location is spatially encoded using DNA barcoding could enrich drug screening and gene expression data; for example, understanding transcriptomic changes in the context of drug penetration. While *in vivo* animal studies are more biologically relevant than two-dimensional (2D) monolayer cell cultures for drug screening^5^, animal models can be costly, time-consuming,^6^ and often fail to translate to humans^7–9^. As an attractive 3D intermediary between homogeneous 2D cell cultures and highly complex tissue samples, 3D cultures have been successfully implemented in cancer and tumour research, and drug screening applications^10^. Recently, organ-on-a-chip platforms have demonstrated the power of 3D cultures to accelerate animal-free clinical drug approvals. Organs-on-chips are microfluidic cultures that recreate organ-level functions and can be used to predict human responses to clinical therapies more accurately than animal models^11^, but despite their significant advantages, organs-on-chips have low throughput and are impractical for use in studies that require many replicates, for example in drug screening and validation^11^. Spheroids, on the other hand, are more readily scalable 3D cell cultures that form sphere-like masses *via* cell-to-cell interactions, which may adopt a structural and spatial complexity similar to that found *in vivo*^12^. 3D cultures can more faithfully represent the heterogeneous cellular responses found in animal models, reflecting gradients in oxygen and nutrient supply, metabolic rate, and cell proliferation. Spatially-resolved analysis of 3D cultures will help to explore sources of heterogeneity in cellular response to external stimuli, including drug exposure.

Characterising gene expression in 3D requires spatial transcriptomics to fully capture variation in gene expression and a complete understanding of the effects of cell-to-cell organisation^13^. Standard single-cell sequencing eliminates spatial information during tissue dissociation, but current spatial transcriptomics techniques can generate spatially-resolved scRNA-seq data^14^. 10x Genomics^15,16^ and NanoString Technologies^17^ offer spatial gene expression platforms whereby tissue sections are mounted on slides with known barcode sequences laid out in an array; capture of transcripts and the known barcode sequence layout on the slide allow gene expression to be mapped to the array, and, by extension, to the tissue. However, these methods currently cannot capture the gene expression profiles of individual cells, since the spatial resolution is currently limited to ∼50 μm^17,18^. In the context of 3D culture models, which are typically only a few hundred microns in size, existing spatial transcriptomic platforms do not have sufficient resolution or high enough RNA capture efficiency to achieve statistically robust sampling of gene expression. Furthermore, the low throughput of current spatial technologies limits analysis of 3D architectures^19^. A method for the study of gene expression that captures profiles at the single-cell level while also maintaining critical spatial information will help to bridge the divide between homogeneous 2D cell culture and animal models.

Here, we construct multi-layered spheroids in which short barcode oligonucleotides (SBOs) encode spatial information and facilitate sample multiplexing for scRNA-seq. We adopt our scTECH-seq method, using SBO–fluorophore conjugates to enable visualisation of barcoded cells in 3D spheroid cultures, prior to scRNA-seq. We demonstrate the use of our platform by constructing HeLa spheroids through a layer-by-layer assembly process and investigating variation in cellular response upon drug exposure. As an example, we profile change induced by the prodrug irinotecan, identifying transcriptional responses that are spatially dependent within 3D cell cultures. We use of fluorescent barcodes also supports correlative analysis, which we demonstrate using a recent mechano-microscopy technique, measuring tissue elasticity within 3D cultures.

## Results and discussion

### scTECH-seq enables visualisation and single-cell sequencing of multi-layered spheroids

scTECH-seq uses short barcode oligonucleotides (SBOs) as described in the 10x Genomics 5′ Gene expression (GEX) kit with library preparation facilitated by the 10x Genomics Chromium platform. Here, we modified the SBOs for the addition of a fluorophore label at the 5′ end, enabling SBO visualisation and quantification in cells (Supplementary Figure S1, Supplementary Table S1). One of the principal benefits of scTECH-seq is the ability to deliver SBOs inside cells, rather than targeting the plasma membrane. Complete internalisation of SBOs helps to limit unwanted exchange between cells and thus allows barcoded cells to be placed back into culture. HeLa cells were transfected for 4 h with dye-conjugated-SBOs (Fig. 1a) using 23 pmol SBOs per 150,000 cells. SBO uptake was visualised with fluorescence microscopy (Fig. 1b), and flow cytometry quantification of fluorescent SBOs indicated >99.9% labelling efficiency (Fig. 1c). A comparison of single-cell sequencing using fluorophore-modified SBOs compared to unmodified SBOs showed no reduction in RNA transcripts per cell (Supplementary Table S2), gene counts (Supplementary Table S3), or SBO numbers per cell (Supplementary Table S4) when using Cy5-modified SBOs compared to unmodified SBOs as per the 10x Genomics standard design (Fig. 1d,e). Having established that fluorophore-modified SBOs did not interfere with library preparation or sequencing, we then extended the barcoding concept by using SBO-labelled cells to construct layered spheroids, in which each layer comprised cells transfected with a unique SBO (Fig. 1f).

**Figure 1.**
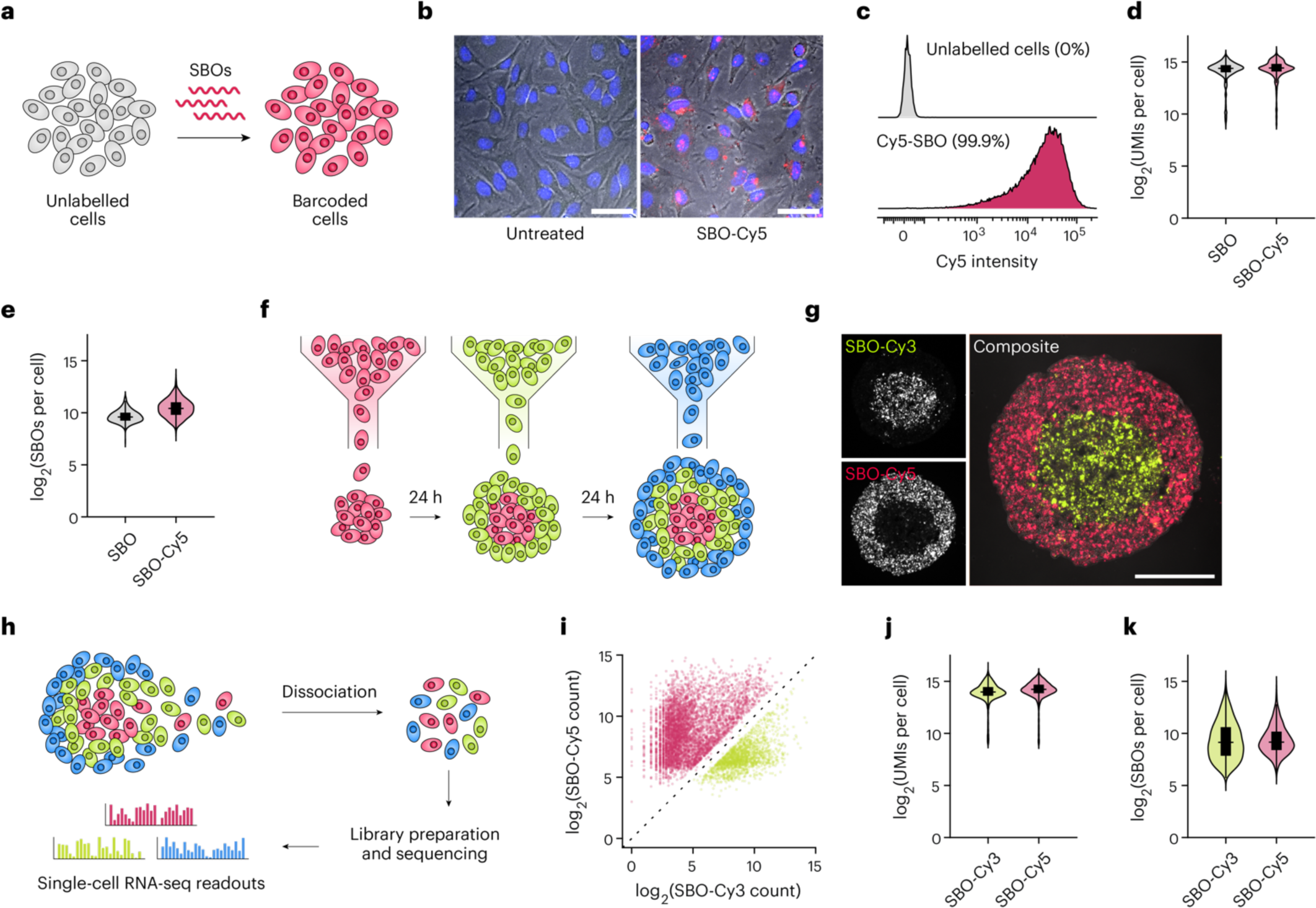
Single-cell transcriptomics of multi-layer spheroids. **(a)** Cells are transfected with SBOs in culture to produce populations of ‘barcoded’ cells. **(b)** Fluorescent microscopy shows SBO (red) uptake in HeLa cells. Scale bar 50 μm. **(c)** SBO uptake efficiency was measured using flow cytometry, with >99% cells positive for SBO fluorescence (two biological replicates, *n* = 5). **(d)** Violin plot showing similar number of RNA counts per cell for SBOs with or without Cy5 modification. **(e)** Violin plot shows similar number of SBO counts per cell with or without Cy5 modification. **(f)** Schematic of the experimental approach for barcoding cell populations and growing spheroids using a layered approach. Different colours represent unique SBO sequences, each encoding a spatial region of interest. **(g)** Representative image of spheroid cross-section showing two cell layers, labelled with Cy5- (red) and Cy3- (yellow) conjugated SBOs. Scale bar 100 μm. **(h)** Schematic of sequencing preparation process. Spheroids are dissociated into single cell suspension and undergo encapsulation and reverse transcription using the 10x Chromium Controller. Data is retrieved and analysed using CellRanger and Seurat. **(i)** Cy5- and Cy3-modifications do not interfere with SBO sequencing and deconvolution using D-score. **(j)** Violin plot shows a similar number of RNA counts per cell for Cy5-SBO and Cy3-SBO. **(k)** Violin plot shows a similar number of SBO counts per cell for Cy5-SBO and Cy3-SBO.

Confocal microscopy of spheroid cryosections enabled clear visualisation of two distinct barcoded layers (Fig. 1g). Subsequent dissociation of spheroids (Fig. 1h) using TrypLE Express and quantification of SBOs *via* flow cytometry confirmed that barcodes were not lost or exchanged between individual cells during the spheroid formation or dissociation process (Supplementary Fig. S2). To confirm that the choice of fluorophore conjugated to SBOs did not disrupt library preparation and sequencing, two-layer spheroids tagged with Cy3- and Cy5-SBOs were dissociated, pooled, and sequenced, whereby the SBOs also facilitated demultiplexing of scRNA-seq data. For each layer, we observed a median of 16,148 and 19,213 RNA transcripts per cell (Fig. 1j, Supplementary Table S5), 4409 and 4884 genes per cell (Supplementary Table S6), and 564 and 571 SBO reads per cell (Fig. 1k, Supplementary Table S7), respectively. Taken together, these results demonstrate a promising capacity to construct and visualise SBO-labelled architectures using unmodified or fluorophore-labelled SBOs without compromising single-cell sequencing reads or demultiplexing integrity.

Next, we analysed the single-cell RNA-seq data obtained from two-layer spheroids. Unsupervised uniform manifold approximation and projection (UMAP) clustering revealed three cell subpopulations based on gene expression that were uniformly distributed throughout the two-layer spheroids (Supplementary Fig. S3). Two of these clusters differed mostly in expression of genes related to different stages in the cell cycle (Supplementary Fig. S4), while the third and smallest subpopulation displayed substantially different expression and strong downregulation of housekeeping and other genes, indicative of potential loss of homeostasis and cell death. We speculate that this minor subpopulation of cells arose during dissociation of spheroids, since cells from both layers of the spheroid were equally represented. Between the two different layers of the spheroids, only 16 genes were differentially expressed (|log_2_FC| > 0.25, *p*_adj_ < 0.05, Supplementary Table S8), representing the first example of spatially dependent gene expression within this architecture. Pathway analysis based on this small set of genes highlighted changes in glucose synthesis and metabolism, CO_2_ accumulation and hypoxia, and some p53-mediated pathways (Supplementary Fig. S5).

### Barcoding facilitates examination of spatial transcriptional differences in multi-layer spheroids

Extending the approach described above, we constructed spheroids from three populations of HeLa cells, each of which had been labelled with a unique SBO. Wree-layer spheroids were grown to a similar size as the two-layer spheroids by adding barcoded cell populations each day over a 3 d period (Fig. 2a). Spheroids were dissociated and sequenced in the same manner as above, with transcript, gene, and SBO counts summarised in Supplementary Tables S9–S11. Using the D-score tool for demultiplexing scRNA-seq data,^4^ 82.5% of cells were unambiguously assigned to one of the three SBOs used, deconvolving cell populations by SBO and hence also by spatial organisation. Cells were clustered into four groups by gene expression during analysis (Fig. 2b). Like the two-layer spheroids, many of the differentially expressed genes between clusters could be attributed to cell cycle progression, and analysis of the cells in clusters 1–3 confirmed substantial differences in the number of cells in G1, S, and G2/M phase (Fig. 2c). Once again, a subpopulation of cells (cluster 4, representing 7.0% of total cells) displayed hallmarks of cell death, including deregulation of housekeeping genes including *ACTB*, *GAPDH*, *HPRT1*, and *EIF* family members. Wese cells were excluded from further consideration.

**Figure 2.**
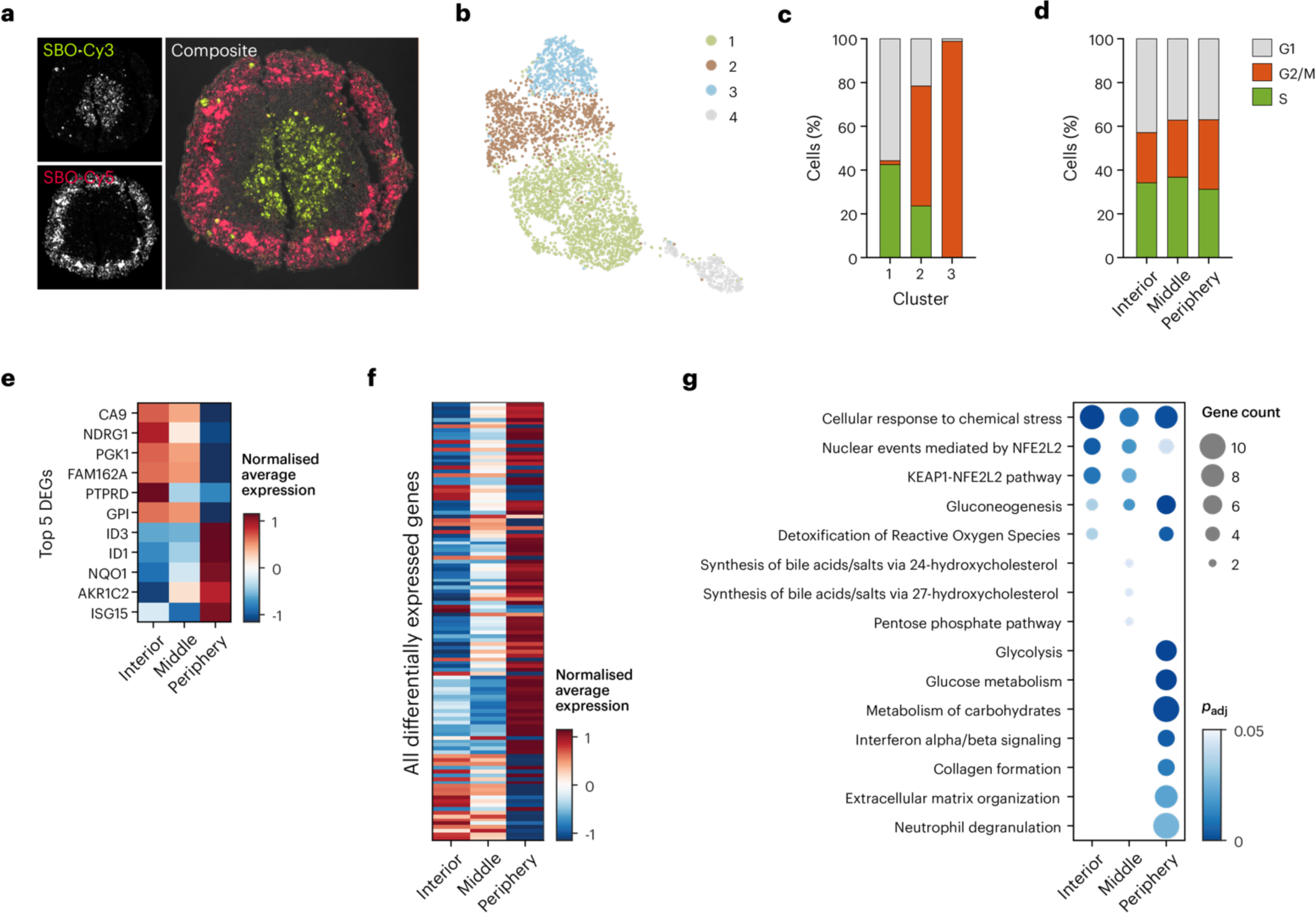
Multi-layer spheroids highlight spatial differences in gene expression related to cell cycle, metabolism, and nutrient supply. **(a)** Representative image of three-layer spheroid. Cells within the core (first layer) are labelled with Cy3-SBO (yellow), the middle layer contains cells barcoded with an SBO that was not conjugated to a fluorophore, and cells on the periphery are labelled with Cy5-SBO (red). **(b)** UMAP representation of the single cells recovered from three-layer spheroids. Cells are coloured by gene expression cluster. **(c)** Quantification of cell cycle markers within each of the three main clusters identified in (b). Cluster 3 principally contains cells undergoing mitosis (G2/M), while cluster 1 displays virtually no markers of G2/M phase. bus, the main differences in gene expression across untreated spheroids are simply due to cell cycle. **(d)** Distribution of cell cycle markers according to spheroid layer show that cell cycle progression is relatively consistent between each of the three layers. **(e)** Normalised average expression levels of the top 5 differentially expressed genes (DEGs, |log_2_ fold change| > 0.25, *p*_adj_ < 0.05) across three-layer spheroids. **(f)** Normalised average expression levels for all differentially expressed genes (|log_2_ fold change| > 0.25, *p*_adj_ < 0.05) across three-layer spheroids. **(g)** Dot plot showing gene ontology (biological process) pathways enriched within each layer of three-layer spheroids. be dot size indicates the gene counts expressed in each pathway, while colour indicates adjusted *p*-value.

Spatial analysis of gene expression revealed that 117 genes were differentially expressed (|log_2_ fold change| > 0.25, *p*_adj_ < 0.05) between the interior, middle and peripheral layers of three-layer spheroids. Examining cell cycle with respect to each of the three layers within the spheroids rather than cluster showed a similar distribution of cell cycle markers across the layers (Fig. 2d), while further examination of the top differential expressed genes across the three layers revealed markers of growth, metabolism, and waste with similarity to those observed across two-layer spheroids (Fig. 2e). We concluded that despite some indicators of cell stress and death within a small subpopulation of cells, cell cycle markers were uniformly distributed across each of the three layers, and the principal transcriptomic differences in multilayer spheroids were simply the result of growth and cell division. Assessment of expression profiles in spheroid interior regions confirmed enrichment of apoptotic and necrotic pathways (Supporting Information Fig. S6).

We top 11 differentially expressed genes (|log_2_ fold change| > 0.25, *p*_adj_ < 0.05) between interior, middle, and peripheral layers in three-layer spheroids are shown in Fig. 2e. 8 of the genes identified as differentially expressed in three-layer spheroids were also differentially expressed in the two-layer spheroids. Qualitatively, of all 117 genes that were differentially expressed (|log_2_ fold change| > 0.25, *p*_adj_ < 0.05) between the three layers, the middle layer tended to display expression levels that were intermediate between the interior and periphery (Fig. 2f), supporting our hypothesis that gene expression patterns could be spatially defined. To investigate whether biological processes differed between spheroid layers, pathway analysis using the differentially expressed genes was conducted (Fig. 2g). KEAP1-NFE2L2 pathways and NFE2L2 nuclear events were more upregulated in the interior, consistent with an increase in oxidative stress, while glycolysis, glucose metabolism, and ECM formation and organisation processes were more prevalent in the peripheral layer than the interior. Wese results are consistent with the current understanding of spheroid cultures in which there is a gradient in metabolic rate, nutrient supply, and oxygen dependent on cell depth, and dense cell-to-cell contact and suppressed cell proliferation in the spheroid core.

Within the three-layer spheroid dataset, we noticed reduced numbers of demultiplexed cells in the interior and middle populations despite our efforts to balance cell numbers during assembly. Since the number of demultiplexed cells correlated with the median number of SBO counts per cell (Supplementary Table S11), we concluded that some barcode degradation had taken place over the longer time taken to grow these multi-layer spheroids, and that there was reduced efficiency of demultiplexing cells from the interior and middle layers. While some SBO degradation in the cellular environment was anticipated, median SBO counts per cell (interior: 564, periphery: 571) were very similar in the two-layer spheroids, suggesting that SBO integrity may be compromised ∼72 h after SBO transfection in HeLa cells.

### *In vitro* analysis of irinotecan-treated spheroids

To explore the potential for spatial analysis of gene expression in response to an experimental stimulus, three-layer spheroids were treated with irinotecan. Irinotecan is a prodrug that is used both alone or in combination with other drugs or treatment options, such as surgery and radiation, for many cancers^20–23^, particularly colon cancer. We primary target of irinotecan is DNA topoisomerase I, an enzyme which is responsible for relaxing supercoiled DNA during replication and transcription^24,25^. Carboxylesterases metabolise irinotecan to ethyl-10-hydroxycamptothecin (SN-38), which as the active metabolite, is several orders of magnitude more potent than irinotecan itself^26,27^. SN-38 stabilises topoisomerase I– DNA complexes, resulting in double stranded DNA breaks and activation of apoptotic pathways^25^. Of note, SN-38 is a fluorescent metabolite with excitation at 360 nm and emission at 420 nm, making it an ideal candidate to monitor drug penetration of spheroids. To determine what concentration of irinotecan to use, we constructed spheroids with three layers and treated them with one of four concentrations of irinotecan prepared in 0.1% v/v DMSO for 24 h, as well as a 0.1% v/v DMSO control (hereafter referred to as ‘vehicle control’) and an untreated control (Supplementary Fig. S7). We percentage of live cells was quantified via flow cytometry using propidium iodide staining as an indicator of viability (Fig. 3a). For subsequent studies, the concentration nearest the IC_20_ (50 μM) was selected.

**Figure 3.**
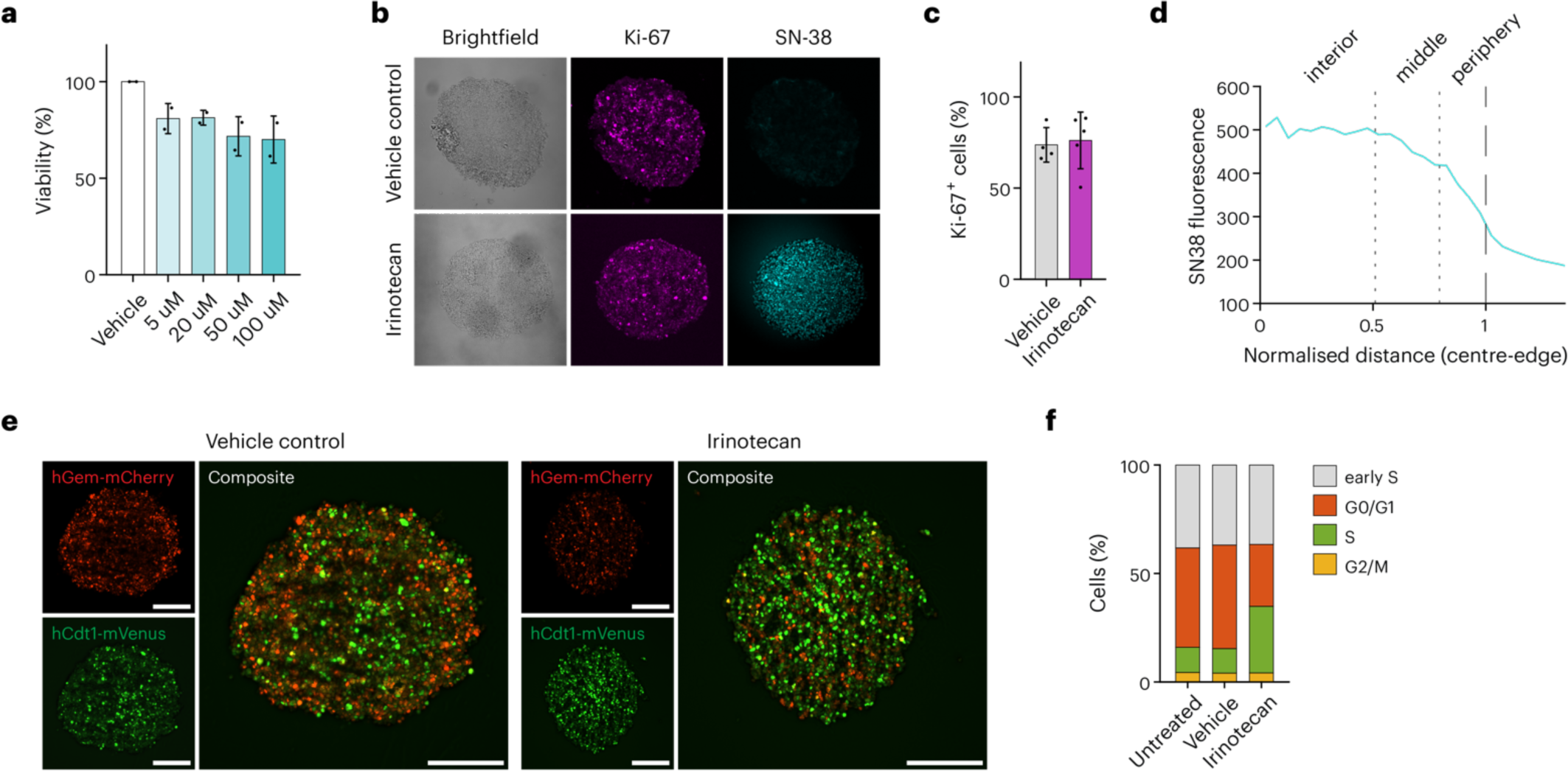
Topoisomerase I inhibition alters cell cycle state in three-layer HeLa spheroids treated with irinotecan for 24 h. **(a)** Percentage of live cells from dissociated spheroids after 24 h irinotecan treatment using propidium iodide. be percentage of live cells (mean ± standard deviation) was calculated via flow cytometry and normalised to the untreated control. 0.1% DMSO (‘vehicle’) is the control. 50,000 single cells were collected per sample. **(b)** Confocal microscopy images of spheroid cryosections stained for Ki-67 detection. **(c)** Percentage of Ki-67^+^ cells from stained spheroid sections (*n* = 4–5 spheroids). **(d)** Quantification of SN-38 fluorescence across the diameter of the spheroid cross-section shown in (c)**. (e)** Representative fluorescence images of Fucci spheroid sections, indicate more cells in S phase. Cells expressing mCherry-hCdt1 (red), mVenus-hGem (green) or both proteins (yellow). mCherry-hCdt1 fluorescence is indicative of G1 phase, mVenus-hGem fluorescence is indicative of S phase, and cells expressing both markers indicate G2/M phase. Scale bar; 200 μm **(f)** Flow cytometry quantification of dissociated Fucci spheroids show an increase of cells in S phase after irinotecan treatment. Spheroids (n = 8) per condition were dissociated and 50,000 single cells were processed during flow cytometry. Flow cytometry graphs are representative of two biological replicates.

As a well-known marker of proliferation in cancer cells, Ki-67 was used to evaluate proliferation in spheroids before (vehicle control) and after irinotecan treatment. Spheroids possessing an apoptotic or necrotic core were expected to have fewer Ki-67^+^ cells within the centre of spheroids, but in this case, immunofluorescence of spheroid sections indicated that Ki-67^+^ cells were distributed homogeneously throughout the spheroids (Fig. 3b). Furthermore, irinotecan treatment did not significantly reduce the number of Ki-67^+^ cells in spheroids when compared to vehicle control (Fig. 3c). Imaging of the fluorescent, active metabolite of irinotecan (SN-38) revealed that the prodrug was metabolised throughout all spheroid layers, as assessed by confocal microscopy (Fig. 3b). Quantification of imaging data indicated the greatest fluorescence intensity within the spheroid core which decreased toward the periphery (Fig. 3d).

Since both irinotecan and its active metabolite SN-38 stabilise topoisomerase I–DNA complexes and induce DNA damage, we assessed cell cycle progression as an indicator of irinotecan activity in three-layer spheroids. To do this, we used fluorescent, ubiquitination-based cell cycle indicator (Fucci) CA technology encoding human Cdt1 (chromatin licencing and DNA replication factor 1 fused to mCherry) and human Geminin fused to mVenus^28,29^. In the Fucci(CA)5 system, cell-cycle stage can be determined by the fluorescence profiles of cells; red fluorescence (mCherry-hCdt1) intensity builds during G2 and remains high through M and G1, dropping sharply in S phase, while green fluorescence (mVenus-hGem) accumulates during S and G2/M, and is rapidly destroyed as cells enter G1. Wus, red fluorescence is indicative of G1, green fluorescence indicates cells in S phase, and the presence of both red and green signals is indicative of cells in late G2/M. Spheroids composed of Fucci-transduced HeLa cells (vehicle control) were sectioned and imaged, revealing a prevalence of mCherry-hCdt1 fluorescence, possibly with some localised cells displaying mVenus-hGem fluorescence in the interior region; in comparison, spheroids treated with 50 µM irinotecan showed an increase in mVenus-hGem expression (Fig. 3e). Wis was corroborated *via* flow cytometry quantification of Fucci markers in dissociated spheroids, showing a 19.4% increase in mVenus-hGem^+^ cells after irinotecan treatment compared to the vehicle control, indicative of cell cycle arrest either prior to completion of S phase or at the G2 checkpoint (Fig. 3f).

### Single-cell RNA sequencing resolves heterogeneity within spheroid layers

Finally, we performed scTECH-seq on HeLa spheroids composed of three uniquely barcoded layers to study the spatial differences in gene expression triggered by irinotecan treatment. Spheroids were treated with 50 μM irinotecan for 24 h before dissociation, library preparation, and single-cell sequencing. After quality control, 11,921 single cells were assigned an SBO using the D-score deconvolution method; 3,748 and 5,674 cells were assigned to the vehicle control and the irinotecan-treated spheroids, respectively. Cells were clustered and a clear separation between the vehicle control and drug-treated cells was observed, consistent with broad changes in gene expression induced by irinotecan treatment (Fig. 4a). A variety of relevant reactome pathways were significantly upregulated in irinotecan-treated spheroids compared to vehicle (Supporting Information Fig. S8).

**Figure 4.**
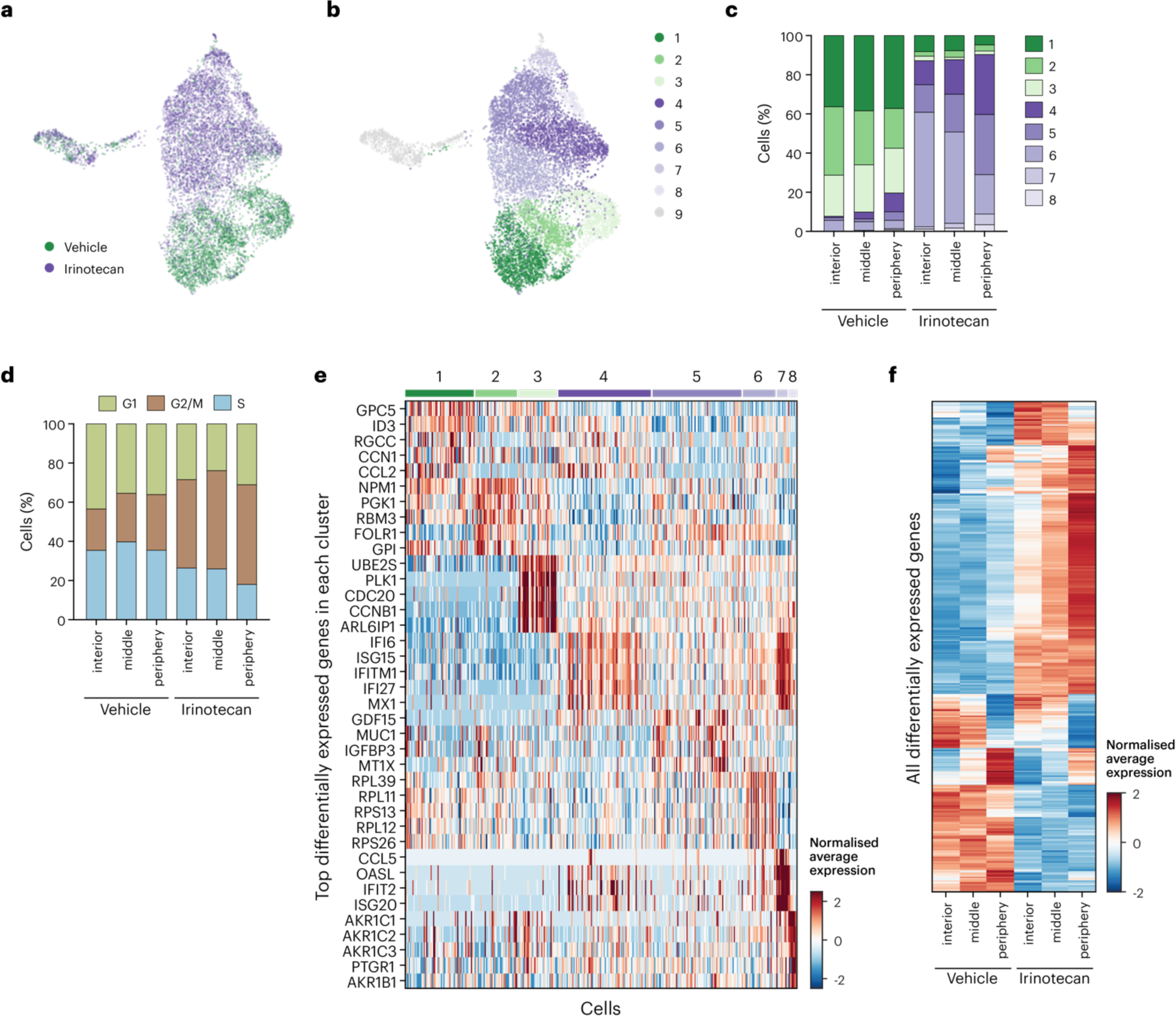
Irinotecan alters transcriptional profiles in multi-layer spheroids. **(a)** UMAP visualisation of single cells captured and sequenced. Cells are annotated according to whether they were treated with 0.1% DMSO (‘vehicle’) or 50 μM irinotecan for 24 h. **(b)** UMAP visualisation of single cells, annotated by cell cluster. **(c)** Distribution of cell clusters according to treatment with vehicle or irinotecan and layer. **(d)** Irinotecan increases expression of G2/M gene markers. Percentage of cells in each cell cycle from the vehicle control and irinotecan treated samples. **(e)** Heatmap enrichment of the top 5 differentially expressed genes (|log_2_ fold change| > 0.25 and *p*_adj_ < 0.05) in each cell cluster. **(f)** Normalised expression levels of differentially expressed genes (|log_2_ fold change| > 0.25 and *p*_adj_ < 0.05) between the vehicle and irinotecan treated spheroids, and each layer.

We set out to identify the different cell subpopulations present within control and drug-treated three-layer spheroids. UMAP clustering revealed the same control populations as in the untreated spheroids, and an additional five clusters that were related to treatment with irinotecan (Fig. 4b). One subpopulation in the dataset (cluster 9) was enriched in markers of cell death in the same way as for the two- and three-layer control spheroids, and since it contained an equal number of cells from each sample, this cluster was excluded in subsequent analysis. With the exception of cluster 2, clusters were distributed across the different layers of the control spheroids in a relatively uniform way, whereas the distribution of clusters within the irinotecan-treated spheroids displayed greater layer-specific differences (Fig. 4c). As an example, cluster 6 contained more cells from the interior region of the spheroids while the remaining clusters were enriched in cells from the periphery. We analysed the different layers for markers of cell cycle phase and found a general increase in G2/M markers in irinotecan-treated spheroids consistent with arrest at the G2 checkpoint (Fig. 4d).

We identified 3,970 differentially expressed genes between the eight clusters (|log_2_ fold change| > 0.25, *p*_adj_ < 0.05). Single-cell expression across the top 5 differentially expressed genes for each cluster is shown in Fig. 4e. Interestingly, cluster 8 displayed an upregulation in aldo/keto reductase (AKR) genes; within this family of stress-related genes, upregulation of *AKR1C3* in particular has been previously identified as a biomarker for irinotecan resistance in colon cancer cells.^30^ Cluster 8 represents a small population of cells displaying a layer-dependent evolution, which is the first evidence that markers of evolving drug resistance with spatial dependence are detectable in our architecture. Visualisation of the average gene expression across spheroid layers for all differentially expressed genes highlighted a variety of patterns, including an array of genes that were significantly upregulated compared to vehicle control in a spatially-dependent manner across spheroid layers following treatment with irinotecan (Fig. 4f).

### scTECH-seq enables spatial analysis of irinotecan-induced change

We unique benefit of our multiplexing model is the ability to classify cells based on spatial location and map drug response across 3D systems. Hence, we explored the three-layer drug-treated spheroids further to investigate different gene expression patterns in response to irinotecan that were dependent on cell depth within the spheroid. We filtered the average expression data for genes that did not vary in different layers of the control spheroids (to exclude baseline variation due to hypoxia, differences in metabolism, *etc.*). Within this subset, we identified 197 genes whose expression level was affected by irinotecan but did not significantly vary between layers (Supporting Information Fig. S9), as well as 39 genes that presented a spatially dependent response to irinotecan (Fig. 5a). Within the set of spatially-dependent genes, *PARP14* has been associated with poor prognosis in pancreatic cancer^31^, *IFITM2* has been identified as a marker of gastric cancer growth and metastasis^32^, and *BCAP31* has been identified as a predictor of inferior survival in non-small cell lung cancer^33^. We identification of a spatially dependent response in cancer-related genes to irinotecan treatment highlights the potential for our system in assessing expression levels of key predictors of cancer progression and survival.

**Figure 5.**
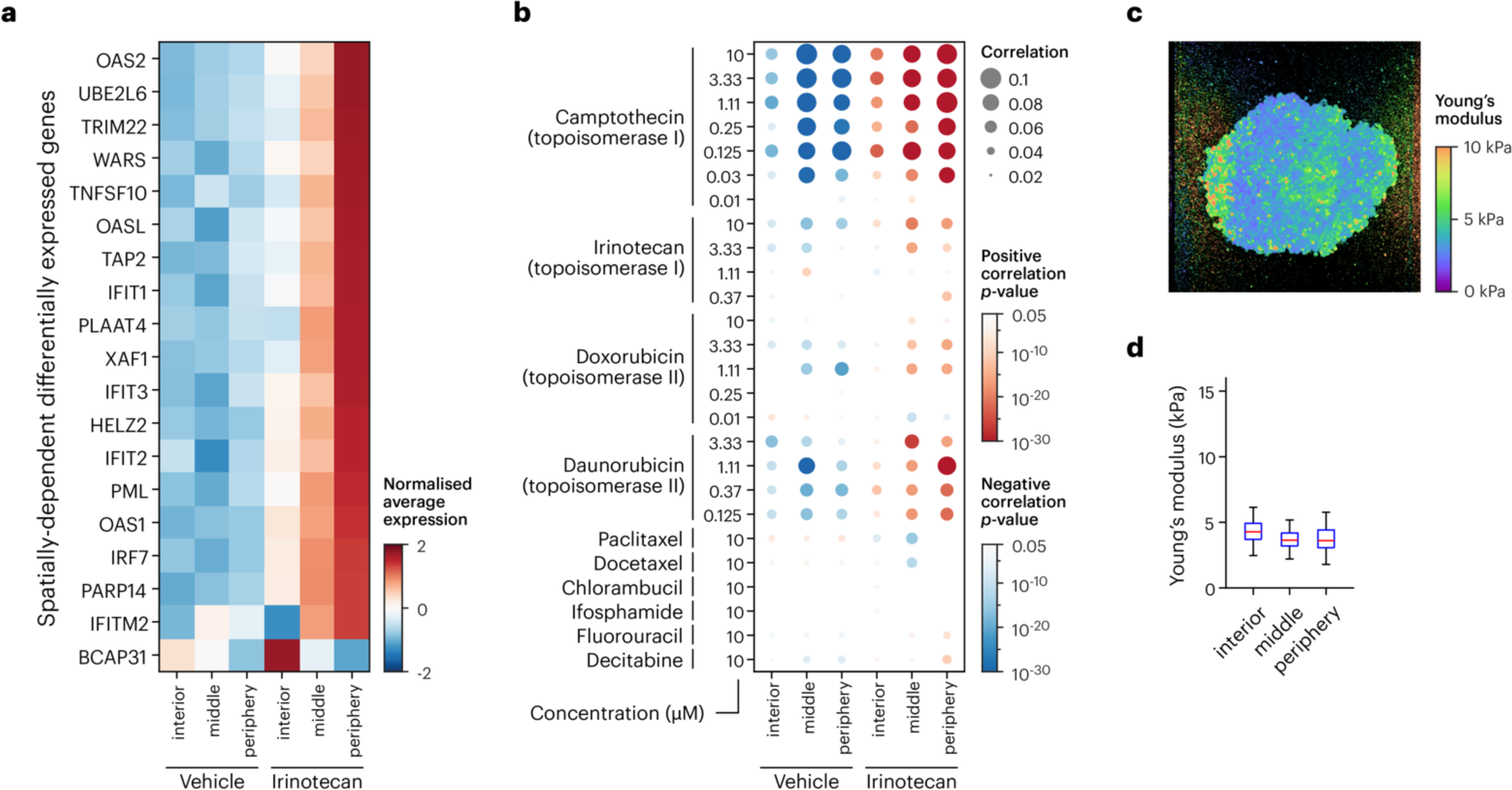
Irinotecan induces spatially dependent gene expression changes in spheroids. **(a)** Normalised average expression levels of differentially expressed genes, filtered for those that display a spatially dependent response to irinotecan while remaining consistent within untreated spheroids. **(b)** Gene expression profiles for each layer in vehicle and irinotecan-treated spheroids were correlated with 24 h HeLa perturbagen profiles from the LINCS L1000 database. Dot size represents Kendall’s rank correlation coefficient τ, red dots show positive correlation, blue dots negative correlation, and colour scales represent *p*-value. DNR: daunorubicin; concentrations shown in µM; drug concentrations 10 µM unless explicitly shown. **(c)** Mechano-microscopy was performed on three-layer spheroids, using fluorophore-tagged SBOs to identify boundaries between layers; the resulting map of Young’s modulus shows that elasticity is relatively uniform throughout three-layer spheroids. **(d)** Spheroid elasticity quantified from the image in (c) did not significantly vary with depth, suggesting that gene expression differences between layers were not induced by variation in mechanical properties.

We next correlated gene expression profiles from each layer of the vehicle and irinotecan-treated spheroids with gene expression profiles from the LINCS L1000 chemical response atlas (Fig. 5b).^34^ LINCS datasets that we employed were all acquired in HeLa cells following 24 h drug treatment, which matches the conditions used in our study. Correlation was based on the average log_2_ fold change calculated for each gene within each of the three layers of the vehicle and irinotecan-treated spheroids, compared to L1000 expression signatures through Kendall’s rank correlation coefficient τ. We observed that in general, peripheral layers displayed the strongest positive correlations, which decreased in magnitude moving toward the interior. Interestingly, much stronger positive correlations were observed with camptothecin rather than irinotecan, suggesting that the transcriptional response of cells to irinotecan is different in 3D spheroids compared to the 2D monolayer cultures used to establish the LINCS database. Considering our imaging results (Fig. 2b), in which there was no apparent concentration gradient in SN-38 suggesting limited drug penetration, we hypothesise that the different gene expression profile may arise through differences in the spheroid environment, including crowding, suppressed metabolic activity, senescence, and hypoxia, all of which were most pronounced in the spheroid core region. We also compared the expression profiles from our spheroids to other drugs; expecting to see some positive correlation with topoisomerase II inhibitors, we profiled against doxorubicin and daunorubicin, where we indeed found positive (albeit weaker) correlations (Fig. 5b). As controls, we profiled against the taxanes paclitaxel and docetaxel, alkylating agents chlorambucil and ifosphamide, and antimetabolites fluorouracil and decitabine, all of which have substantially different modes of action to topoisomerase inhibitors, and accordingly we found only very weak similarities in gene expression. Our results are consistent with the hypothesis that spatially encoded spheroid architectures can be used to profile differential transcriptional responses to drugs in a 3D context. Furthermore, we find that drug expression profiles may differ between 2D and 3D models, and also with cell depth in spheroidal architectures. Transcriptomic data at the single-cell level will likely be a requirement for establishing future perturbagen atlases in 3D culture models.

Finally, to investigate the source of spatial differences in cellular response to irinotecan, we examined the mechanical properties of spheroids grown in the layer-by-layer approach. Our hypothesis was that there may have been a discontinuous structure at spheroid layer boundaries because of stepwise growth with periods of incubation in between, which could lead to variation in drug penetrance. We examined the mechanical properties of three-layer spheroids using mechano-microscopy, a high-resolution variant of quantitative microelastography. Spheroids were embedded in hydrogels, and the deformation in response to an applied force was measured to obtain elasticity maps. Capitalising on the fluorophore-labelled SBOs present within each of the spheroid layers, we were able to assign elasticity measurements to each of the three regions. We multi-layered spheroids did not exhibit any signs of a disjoint boundary or change in elasticity between the layers (Fig. 5d). Wus, we conclude that the spheroids were relatively homogeneous in structure, even though they were assembled in three distinct stages, and that the variation in response to irinotecan was likely due to differences in drug penetration or cellular metabolic activity and not microenvironment. Interestingly, in comparison to our layer-by-layer assemblies, spheroids of a similar size grown in a single step displayed an elasticity that varied inversely with the cellular depth (Supplementary Fig. S10).

## Discussion

In this work, we have demonstrated the ability to conjugate fluorophores to short barcode oligonucleotides (SBOs), which does not affect reverse transcription, single-cell library preparation, or sequencing. Our transfection-based, antigen-independent cell barcoding method (scTECH-seq) uses polymer-based transfection, removing the requirement for a highly labile surface-based antibody or lipid labelling approach. While polymer transfection is still a transient labelling technique, the SBO lifetime is sufficient to allow transfection, cell harvesting, and subsequent assembly and treatment of novel 3D architectures over a period of 72–96 h. We found evidence of SBO degradation that would likely compromise successful demultiplexing after this timepoint, and it should be noted that the persistence of fluorophore signals within cells is not necessarily indicative of SBO sequence integrity.

However, since polymer-based delivery of SBOs is known to result in endolysosomal entrapment, exploring alternative delivery reagents to target alternative uptake routes^35^, modifying SBOs with phosphorothioate linkages to reduce nuclease-mediated degradation^36^, or generating stable cell lines with genetically encoded barcode sequences and/or fluorescent reporters^37^ are all strategies that could help to preserve SBO integrity over longer time periods in future work.

We addition of fluorophores to SBOs serves several purposes: (i) it enables visualisation of SBO^+^ populations, (ii) it can be used to accurately quantify SBO uptake, and (iii) it can facilitate FACS to selectively remove cell subpopulations containing multiple SBOs prior to library preparation. We anticipate that SBOs modified in this manner may enable cell sorting based on SBO identity, for example, to clean up populations for cell types with non-uniform SBO uptake efficiency, or prior to sequencing to remove cells containing non-unique SBOs. We further envision that assembly approaches could be extended beyond our layer-by-layer approach to automated 3D bioprinting to increase throughput and reproducibility.

While spatial transcriptomics platforms are ideal for exploring gene expression over comparatively large tissue sections, spatial transcriptomics for spheroids (even as large as ours, several hundred µm in diameter) is limited by the resolution of capture spots, which is generally 50–220 µm with a centre-to-centre gap of 100-500 µm. Many spheroid sections would need to be analysed and the data manually combined to provide statistically robust data, and even then, the resolution would be poor. A previously reported technique for spatial transcriptomics in 3D cultures makes use of dye penetration to indirectly quantify cell depth in 3D cultures prior to single-cell analysis^38^. However, our technique is not limited to measurements of diffusion and could be used to profile aspects of 3D culture, including proliferation or epithelial–mesenchymal transition, which may not be directly related to diffusivity, especially in more complex architectures.

As we have demonstrated, SBOs not only facilitate multiplex single-cell RNA-seq, but can also encode the spatial arrangement of cells, allowing single-cell transcriptomic data to be related to cell location within engineered 3D cell culture architectures. Using a stepwise growth of multi-layered spheroids, whereby cells in each layer contain a unique SBO, we performed multiplex single-cell RNA sequencing to map spatial differences in cellular response within 3D cultures. Our data support the hypothesis that spheroid cultures display limited cell growth and proliferation within the core due to dense cell packing and a limited supply of nutrients and oxygen. In some cases this can also result in a necrotic core^39,40^. Exploration of cellular response to irinotecan in the 3D context revealed both uniform changes in gene expression throughout the entire spheroid, as well as genes that displayed expression level gradients in response to drug exposure. We also observed upregulation of potential indicators of drug resistance evolving in a spatially dependent manner, particularly within cells at the periphery of spheroids, presumably due to exposure to a higher drug concentration. While our analysis by imaging SN-38, the active metabolite of irinotecan, was consistent with a previous study that quantitatively demonstrated SN-38 accumulation in the core of spheroids using MALDI-MS^41^, our sequencing results suggest a gradient in the transcriptomic response that indicated the greatest drug concentration and drug-related effects at the periphery. Extending the fluorescent barcoding idea further, the visualisation of SBOs within spheroids also supported *in situ* correlative imaging techniques; we demonstrated this possibility using mechano-microscopy to construct maps of elasticity (*i.e.,* Young’s modulus) in 3D cultures. We ability to relate single-cell sequencing data to matrix elasticity could facilitate correlative studies of gene expression in the context of mechanobiology.

In this study, we highlight that transfection-based hashing methods (such as scTECH-seq) are compatible with the stepwise growth of layered spheroids. We modification of SBOs with a fluorophore enables cell subpopulations within the assembly to be imaged, and the use of unique SBO sequences facilitates multiplexing of single-cell RNA sequencing data to collect spatially-resolved, transcriptomic information. Our spatially-resolved data highlights differences in gene expression that could be used to inform the first 3D gene expression atlases in response to drug treatment. Using scTECH-seq, the construction of engineered cellular architectures adds spatial resolution to studies of gene expression using *in vitro* 3D culture models.

## Methods

### SBO Barcode design

We 10x Genomics 5′ feature short barcode oligonucleotides (SBOs) were ordered from Integrated DNA Technologies (IDT Asia Pacific). SBOs (69 nt) were designed according to 10x Genomics specifications, except with the following modifications: oligonucleotides were modified by replacing the C_12_ amine linker with a shorter chain internal modified C (/iAmC6C/) and adding a fluorophore at the 5′ end (/5Cy5/ or /5Cy3/). Oligonucleotides were ordered at 250 nmol scale with HPLC purification. A representative SBO sequence using feature barcode 13 shown in lowercase is 5′-/5Cy3//iAmMC6C/GGAGATGTGTATAAGAGACAGNNNNNNNNNNataatcattacgtggNNNNNNN NNCCCATATAAGA*A*A-3′ (* = phosphorothioate). SBO sequences used in this study are presented in Supplementary Table S1.

### Cell culture

HeLa cells were cultured in Dulbecco’s Modified Eagle Medium (DMEM) (Gibco 10569044) supplemented with 10% v/v FBS at 37 °C and 5% CO_2_. HeLa cells were maintained at 37 °C in a humidified incubator with 5% CO_2_ and passaged at 70–80% confluency using 1× TrypLE Express (Gibco).

### Transfection protocol

Dendritic polymer for transfection of SBOs was synthesised as previously described^42^ using a 25 mol% GMA backbone and generation 5.0 poly(amido amine) dendrons. Polymer and SBO complexes were formed at amine-to-phosphate (N/P) ratio 30. For SBO labelling, cells were seeded at 150,000 cells/mL in a 24-well plate and incubated overnight at 37 °C and 5% CO_2_. Polymer (3 mg/ml stock solution in ddH_2_O) and SBO (100 µM stock solution in ddH_2_O) were diluted in Opti-MEM (Gibco 31985070). SBO was diluted to 657 nM for a total of 23 pmol (1.59 nmol DNA phosphate groups) in 35 μL, while the polymer was diluted to 0.326 mg/ml for a total of 47.7 nmol primary amines in 35 µl. Equal volumes of polymer (34 µl) and SBO (34 µl) solutions were mixed gently but thoroughly and incubated for 25 min at room temperature. Cells were washed with 1×PBS once to remove serum and media was replaced with 150 μL Opti-MEM. 65 μL polymer–SBO complexes were added per well and cells were incubated at 37 °C for 4 h. Transfection efficiency was visualised with an epifluorescence microscope and quantified *via* flow cytometry.

### Flow cytometry

For 2D cell cultures, transfection solution was removed and cells were washed with PBS twice. Cells were collected with TrypLE and 2× FACS buffer (2 mM EDTA + 6% FBS in 1×PBS) for flow cytometry. For spheroid dissociation, cells washed with PBS twice before incubation with TrypLE for 20 min at 37 °C. Spheroids were manually dissociated by pipette mixing 20–30 times until cell clumps were no longer visible. Cells were collected by centrifugation (300*g*, 5 min) and resuspended in 2× FACS buffer.

### Spheroid formation

To form spheroids, transfected HeLa cells were plated in Nunclon Sphera 96-well U-shaped-bottom microplates (Wermo Scientific 174925) and incubated for at least 24 h. Spheroid formation was visualised under phase contrast microscopy. To grow three-layer spheroids, HeLa cells were transfected as described above. Transfected cells were harvested and plated in 96-well low adhesion plates (Nunclon Sphera) at 2,000 cells/well in 100 μL media. To form the second layer, the following day, a new sample of HeLa cells was transfected with a different SBO prior to harvesting and 3,000 cells/well were seeded into fresh wells. Spheroids formed the previous day were transferred to the new well, with one spheroid per well. Wis process was repeated on the third day, using 5,000 cells/well to form the final layer. Each day, cells were incubated for 24 h prior to the addition of the next layer. Two-layer spheroids were prepared the same way using 5,000 cells/well for both layers.

### Spheroid dissociation and viability assay

Propidium iodide was used to assess the proportion of live/dead cells in spheroids after irinotecan treatment. 24 h after the third layer of cells were added, spheroids were treated with irinotecan. Irinotecan hydrochloride (Sigma-Aldrich I1406) was dissolved at 50 mg/mL in DMSO. Irinotecan was diluted in media to a 1 mM working stock and serial dilutions were used to make 200 μM, 100 μM, 40 μM, and 10 μM dilutions. Half of the volume in each well was removed and diluted irinotecan stocks were added in equal volume to achieve final concentrations of 100, 50, 20 and 5 μM. Spheroids were incubated for 24 h. Vehicle (0.1% v/v DMSO) and untreated spheroids were used as controls. After 24 h, spheroids (*n* = 8) for each treatment were transferred into a 1.5 mL tube. Media was removed and 300 μL 1x TrypLE Express was added, and spheroids were incubated for 15 min at 37 °C. After 15 min another 300 μL TrypLE was added to each tube and spheroids were pipetted up and down 20 times using a 1000 µL single-channel pipette until no visible cell clumps remained. Cells were centrifuged at 300*g* for 5 min and resuspended in 2× FACS buffer supplemented with 4 µg/mL propidium iodide solution. Cells were stored on ice until flow cytometry was performed (BD SORP Fortessa), using 561 nm laser and 610/20 nm bandpass filter. Data was analysed using FlowJo v10.

### Spheroid sectioning

Following spheroid growth or drug treatment, spheroids were transferred into 1.5 mL tubes. Media was removed and 4% paraformaldehyde (PFA) in 1×PBS was added and spheroids were incubated for 20 min at 25 °C. Spheroids were washed with 1×PBS twice and incubated in 30% sucrose overnight at 4 °C for cryoprotection. We following day, spheroids were frozen in optimal cutting temperature (OCT) compound for cryosectioning using 12 μm slices.

### Ki-67 immunofluorescence

Hydrophobic barrier pens were used to create a water-repellent barrier around spheroid sections. All incubation steps were conducted at 25 °C unless otherwise stated. Spheroid sections were fixed to slides with 4% w/v PFA for 15 min and washed with 1×PBS twice, 5 min each. Sections were permeabilised with 0.1% v/v Triton X-100 in 1×PBS for 10 min, followed by 2 × 5 min washes in 1× PBS. Blocking was conducted using 2% w/v BSA in 1×PBS for 1 h and incubated with anti-Ki67 antibody (Abcam ab16667) in a 1:200 dilution, overnight at 4 °C. Sections were washed 3 × 5 min in 0.1% v/v Tween 20 in 1×PBS. Secondary antibody incubation was left for 1 h using anti-rabbit Alexa Fluor 488-conjugated at a 1:500 dilution. Sections were washed with 0.1% v/v Tween 20 in 1×PBS twice and 1×PBS before mounting glass coverslips with Fluoromount G (Southern Biotech). Images were acquired with a Nikon A1 RMP confocal using a 10× objective and analysed using ImageJ.

### Fucci cells

Stable HeLa-Fucci cell line was generated using the Super PiggyBac Transposase Expression Vector system (System Biosciences, PB210PA-1) together with tFucci(CA)5 plasmid^29^ (Addgene #153521), as per manufacture’s protocol. Briefly, HeLa cells were seeded at a density of 80,000 cells/mL in a 12-well plate and left to settle overnight (∼16 h). For transfection, 200 ng of the PiggyBac transposase was mixed with 500 ng of tFucci(CA)5 plasmid (overall ratio of 1:2.5 transposase: transposon vector), and diluted to a total of 30 µL in Opti-MEM (Gibco 31985070). Separately, 0.7 µL Lipofectamine 2000 (Invitrogen 11668027) was diluted to 30 µL in Opti-MEM, and the two solutions were mixed thoroughly (total of 60 µL transfection cocktail) and incubated for 25 min at r.t. In the meantime, cell media was removed, cells were rinsed twice with 1×PBS, and 400 µL Opti-MEM was added. After incubation, 60 µL of the transfection cocktail was added to the well, and cells were incubated at 37 °C for 3 h, at which point 540 µL of DMEM + 10% v/v FBS media was added to the well and the transfection was left to proceed for a further 45 h (48 h total). After 48 h, the cells were transferred to a 10 cm plate and left overnight to settle. Cells were routinely cultured in selection media (DMEM, 10% FBS, Blasticidin S HCl 10-30 µg/mL) until 100% of cells expressed the protein markers.

### Single-cell sample preparation and sequencing

Wree-layer spheroids were formed as described above. Spheroids were treated with 50 μM irinotecan or 0.1% DMSO for 24 h. Spheroids were dissociated as outlined in ‘Spheroid dissociation and viability assay’. Single cells from the vehicle control and irinotecan spheroids were counted and pooled in equal number at 2,000 cell/μL in 0.4% w/v BSA for sequencing. Cell viability was confirmed to be >90%. Cells were loaded onto a 10x Genomics Chromium Controller, Chip K (1000286) and libraries were prepared using the Chromium Next GEM single cell 5′ kit v2 and 5′ Feature Barcode kit following manufacturer guidelines (10x Genomics 1000263, 1000265). Illumina iSeq 100 platform was used for initial QC of the libraries and deeper sequencing was conducted on Illumina NovaSeq 6000 in the vendor recommended format to reach up to 50,000 reads per cell. Fastq files were generated with Illumina bcl2fastq v2.20.0.422, allowing 1 mismatch in each index. 10x Genomics Cell Ranger was used for transcript alignment and generation of gene count and feature barcode matrices.

### Single cell data analysis

We D-score deconvolution algorithm was used to assign SBOs to single cells (available at https://github.com/genomicswa/d-score). Wree-layer spheroids are used here as an illustrative example. A total of 11,921 single cells were sequenced and successfully assigned an SBO. Cells were filtered for 200–95,000 total genes and less than 15% mitochondrial genes detected. 9,422 cells were used in downstream analysis. Data were normalised and the top 3,000 variable genes were selected for scaling. We scaled data was used for principal component, dimensional reduction and UMAP clustering. We Wilcoxon rank sum test was used to calculate differentially expressed genes (|log_2_ fold change| > 0.25, *p* < 0.05). Differentially expressed genes were corrected for multiple testing using adjusted *p*-value. We following software versions were used: Cell Ranger v7.0.0, ClusterProfiler v4.6.2, GSEABase v1.60.0, ReactomePA v1.42.0, Seurat v4.3.0.1.

### Mechano-microscopy

Mechano-microscopy was conducted as previously described^43^. Briefly, the mechano-microscopy system is built upon an optical coherence microscopy (OCM) setup using a supercontinuum laser for broadband light. We Michelson interferometer in the OCM system includes filters to create a spectral range of 650 nm to 950 nm and achieves ∼1.4 µm axial resolution and ∼0.5 µm lateral resolution. Two B-scans were acquired per location and B-scans were analysed to measure local displacement and strain, allowing for elasticity estimation using the stress–strain response of a pre-characterised compliant layer placed at the bottom of the sample. A confocal fluorescence microscopy system and a photomultiplier tube detector is integrated alongside the OCM to visualise fluorescent barcodes.

## Data and Code availability

We R code used to generate the plots presented in the main figures are available at https://github.com/Jessica736/Layer-by-layer-spheroids. We single-cell sequencing datasets generated in this study have been deposited in the National Centre for Biotechnology Information (NCBI) Gene Expression Omnibus (GEO) under accession number GSE245416.

## Author information

Corresponding author: Cameron W. Evans.

## Author contributions

J.J.K. performed cell culture, barcoding, and spheroid experiments. M.N. synthesised and characterised dendritic polymers. J.A.K. transduced HeLa cells with Fucci. J.J.K. and S.S.A.M. prepared and imaged spheroid sections. A.M., S.E.A., Y.S.C. and B.F.K. performed elasticity measurements. M.I., U.D.K. and A.S. processed cells for sequencing and performed initial bioinformatics processing of the dataset. J.J.K. and C.W.E. processed and analysed the sequencing data and prepared figures. N.M.S., A.S. and C.W.E. conceived experiments. N.M.S. and C.W.E. supervised the project. N.M.S., K.S.I., and C.W.E. acquired funding. J.J.K. and C.W.E. wrote the draft manuscript. All authors contributed to proofreading and editing the manuscript.

## Conflict of interest statement

None to declare.

## Acknowledgements

Wis work was funded by the Australian Research Council and the State Government of Western Australia through the Future Research & Health Innovation Fund. We acknowledge the facilities and assistance at the Centre for Microscopy, Characterisation and Analysis at the University of Western Australia. Genomics WA is supported by BioPlatforms Australia, the State Government of Western Australia, Harry Perkins Institute of Medical Research, Cancer Research Trust, Telethon Kids Institute and the University of Western Australia. We gratefully acknowledge the Australian Cancer Research Foundation and the Centre for Advanced Cancer Genomics for making available Illumina Sequencers for the use of Genomics WA. We acknowledge the facilities, and the scientific and technical assistance of Microscopy Australia at the Centre for Microscopy, Characterisation & Analysis, We University of Western Australia, a facility funded by the University, State and Commonwealth Governments. J.A.K. acknowledges the Forrest Research Foundation for a Forrest Postdoctoral Fellowship.

## Notes

https://www.ncbi.nlm.nih.gov/geo/query/acc.cgi?acc=GSE245416

### Competing Interest Statement

The authors have declared no competing interest.

