## Supplementary information for "Polymer-mediated oligonucleotide delivery enables construction of barcoded 3D cultures for spatial single-cell analysis"

#### Supplementary Figures

**Supplementary Figure S10.** Layered spheroids display uniform stiffness. **(a)** Optical coherence tomography (OCT) (left) and mechano-microscopy (right) images of mono-layer spheroid. **(b)** Box plot of elasticity measurements for pixels across the three arbitrarily selected regions of interest (R1-3) outlined in (a). Mean and standard deviation (SD) are also shown. **(c)** Fluorescence, OCT and mechano-microscopy images of three-layer spheroid. Cells in the core and outer layers were both tagged with Cy3-SBO to identify boundaries between the different layers since the setup for imaging in conjunction with elastography was a single-colour excitation system. **(d)** Elasticity (Young's modulus) measurements for pixels across the three layers (R1-3) outlined in (c) indicate little change in stiffness between the layers. .......... 11

### Supplementary Tables

### Supplementary Figures

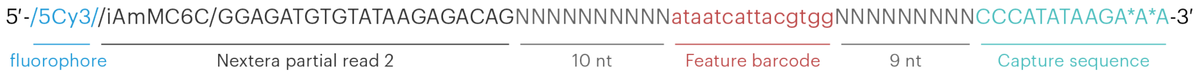

**Supplementary Figure S1.** Short barcode oligo (SBO) design, incorporating fluorophore at the 5′ end.

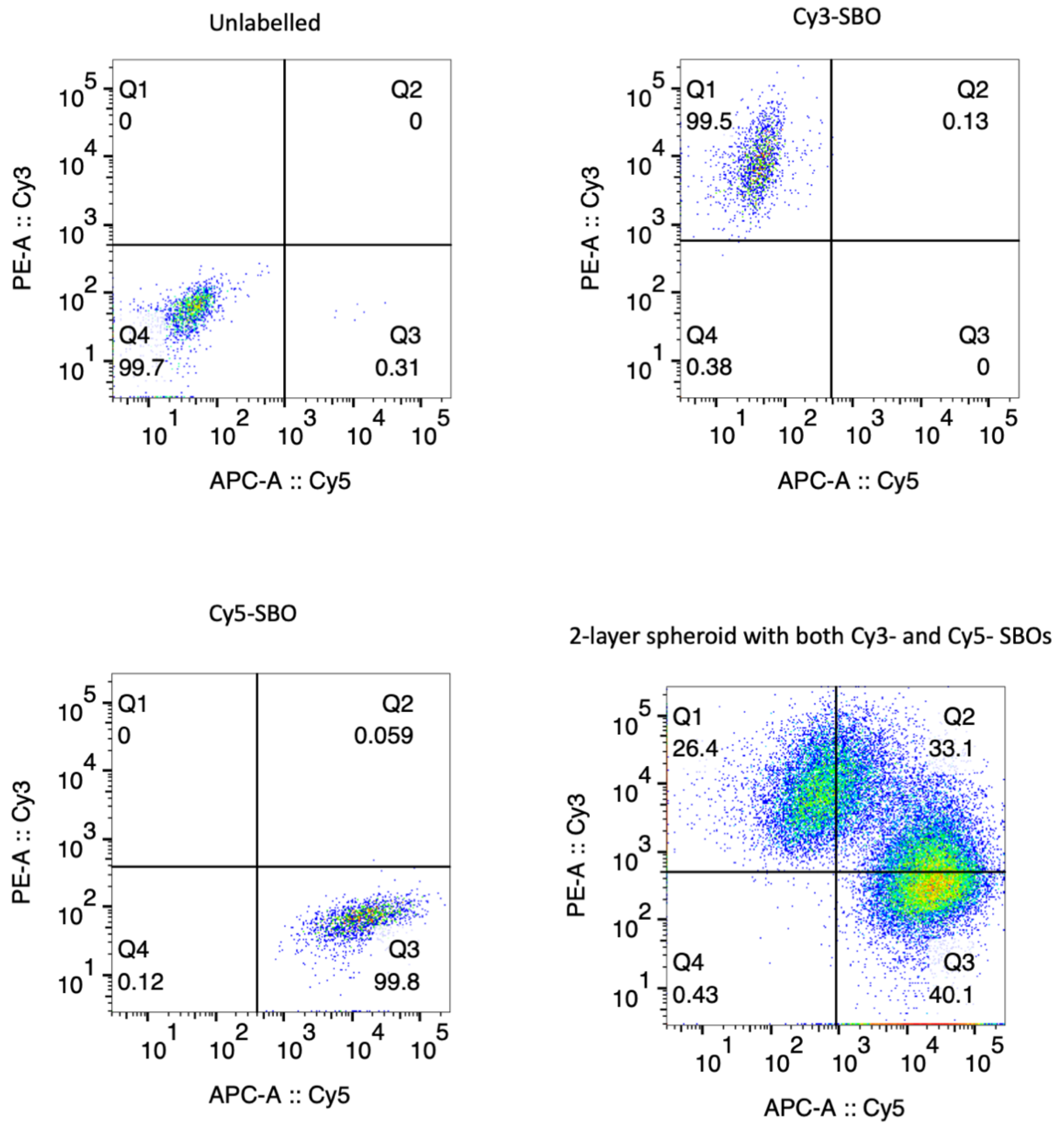

**Supplementary Figure S2.** Cyanine fluorophore-tagged SBOs can be used for sorting cells with a single SBO. Dissociated single cells from two-layer spheroids containing Cy5-SBO and Cy3-SBO labelled cells maintain their SBOs and are still resolved as distinct populations. 80,000 single cells were collected.

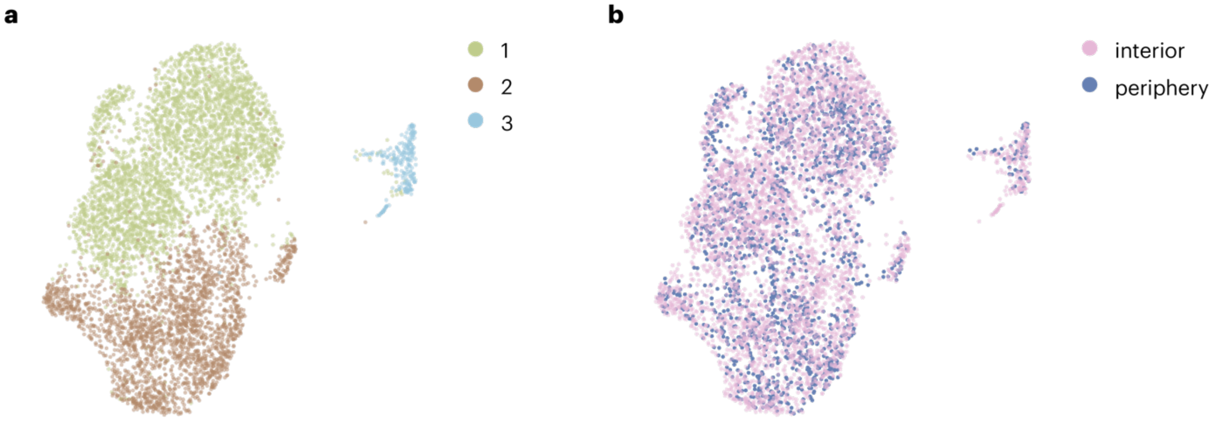

**Supplementary Figure S3.** UMAP in two-layer untreated spheroids. **(a)** Cells are clustered into three subpopulations. **(b)** Cells mapped in the same UMAP plot according to cell location within spheroids shows that gene expression patterns are not spatially dependent since cells from the interior and periphery are found uniformly across all clusters.

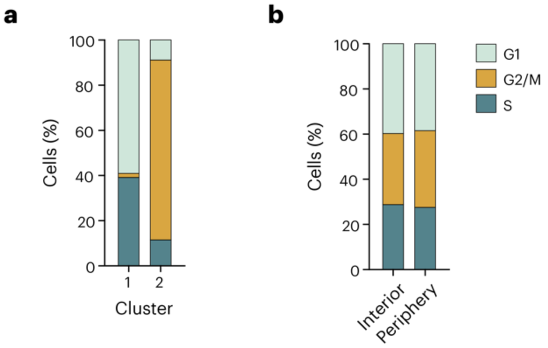

**Supplementary Figure S4.** Cell cycle in two-layer spheroids. **(a)** Cell cycle evaluated by cluster shows the principal differences between the two major clusters is due to mitosis. **(b)** Cell cycle evaluated according to spheroid layer shows that cell cycle is evenly distributed throughout the whole spheroid. These plots are consistent with observations in Supplementary Figure S2.

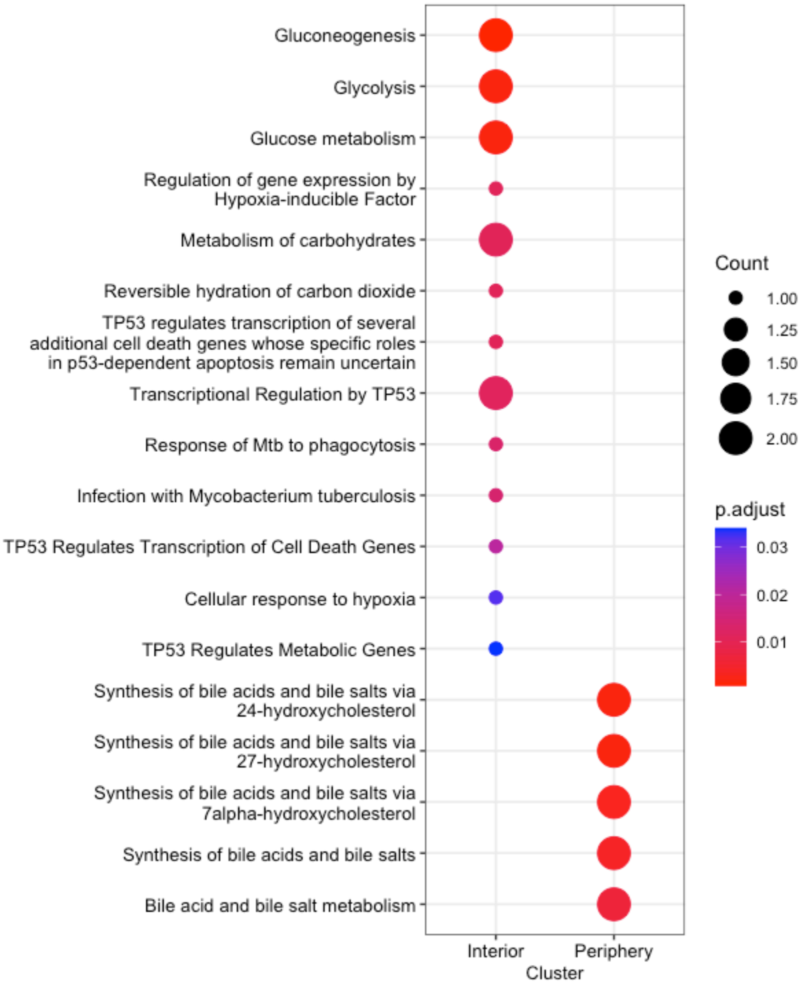

**Supplementary Figure S5.** Pathway analysis in two-layer spheroids.

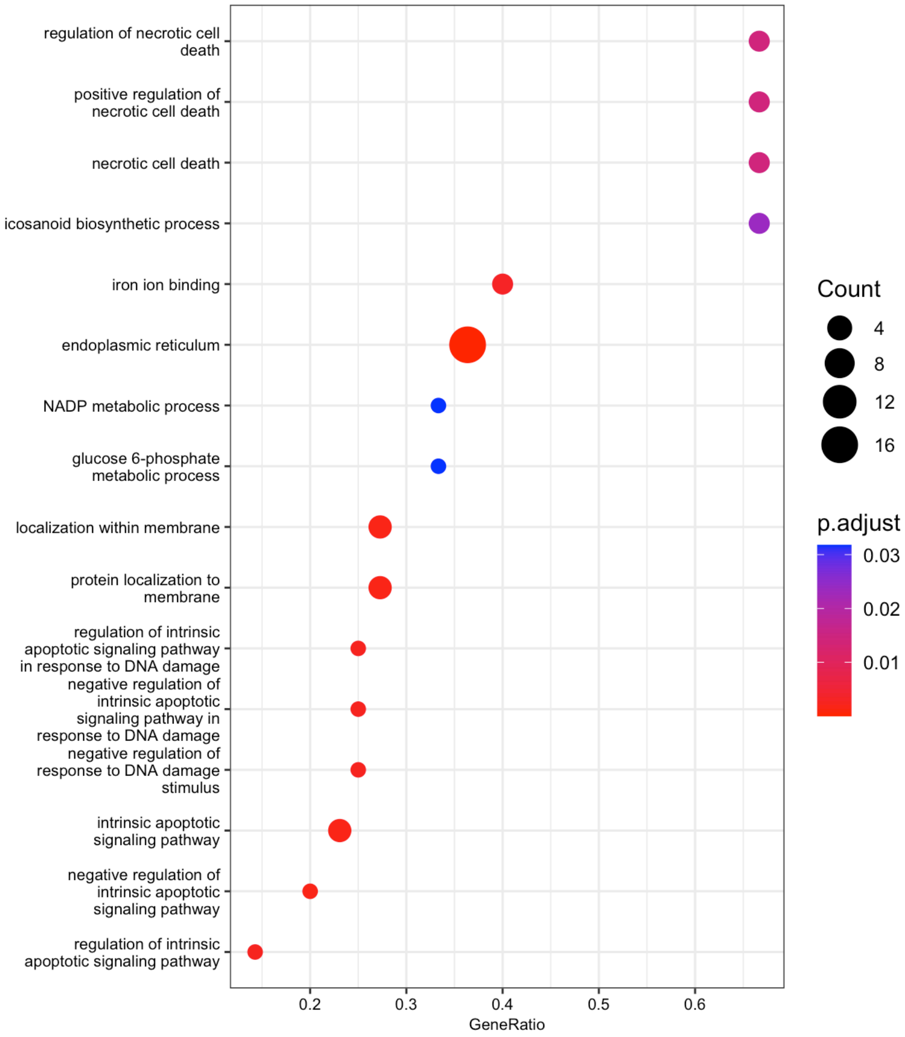

**Supplementary Figure S6.** Gene ontology (biological processes) enrichment demonstrate that the cells within spheroid cores are enriched for apoptotic and necrotic pathways. Dot size indicates gene counts expressed in each pathway and colour indicates adjusted *p*-value.

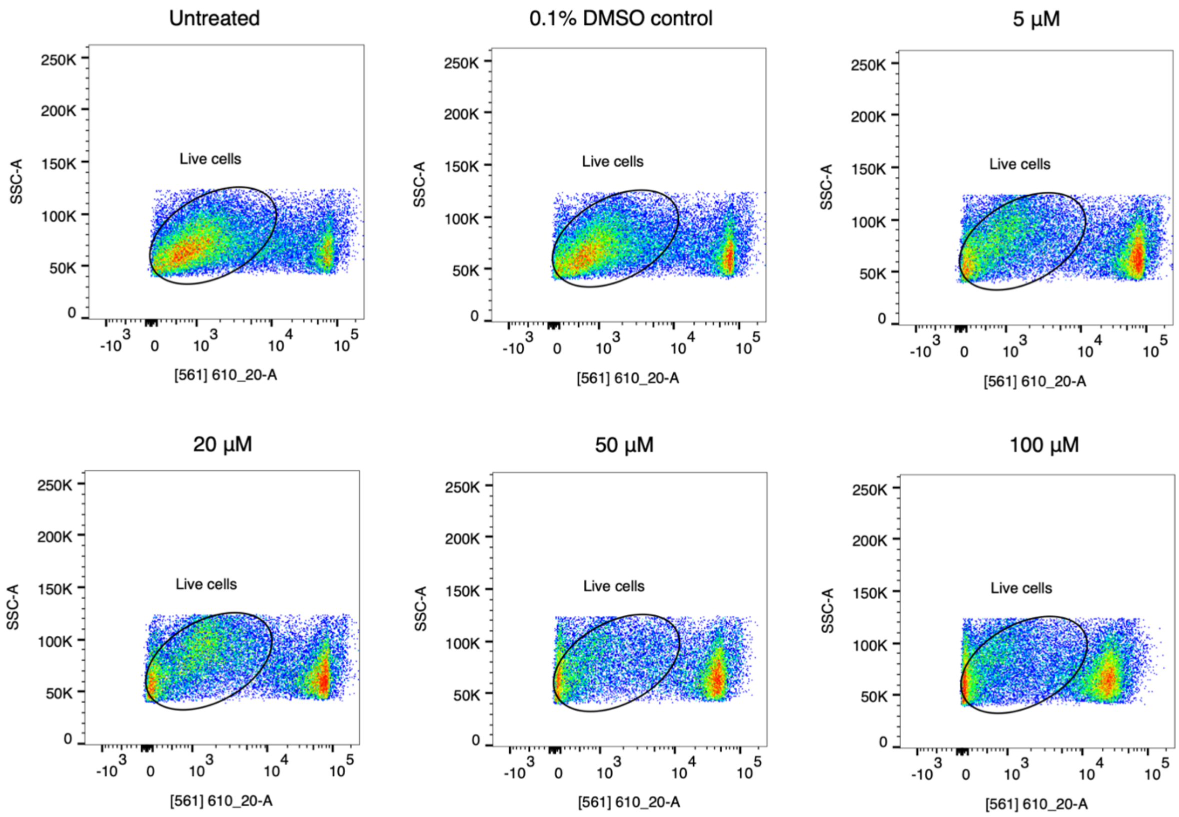

**Supplementary Figure S7.** Flow cytometry plots indicating live cell populations with propidium iodide staining. 30,000 single cells were collected for each sample.

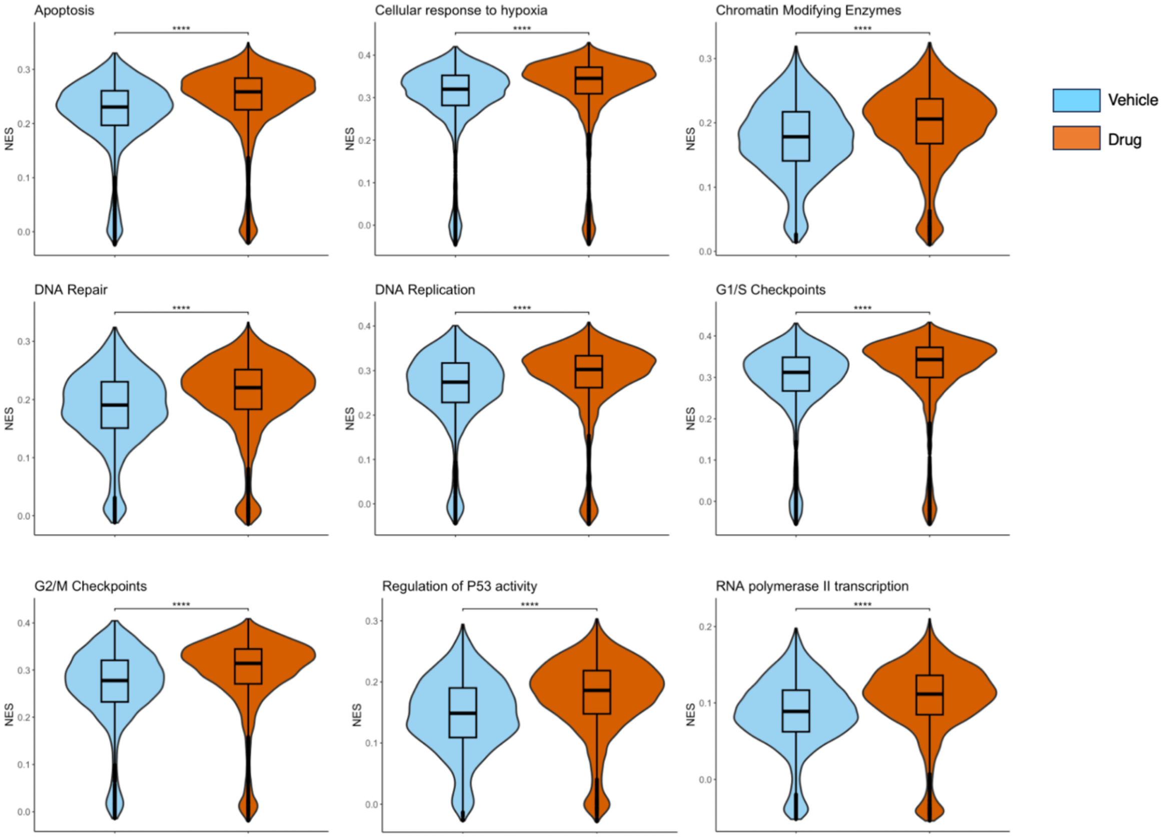

**Supplementary Figure S8.** Normalised enrichment scores for Reactome pathways in response to irinotecan treatment. Statistical significance calculated between vehicle control and irinotecan-treated cells; **** $p < 0.0001$.

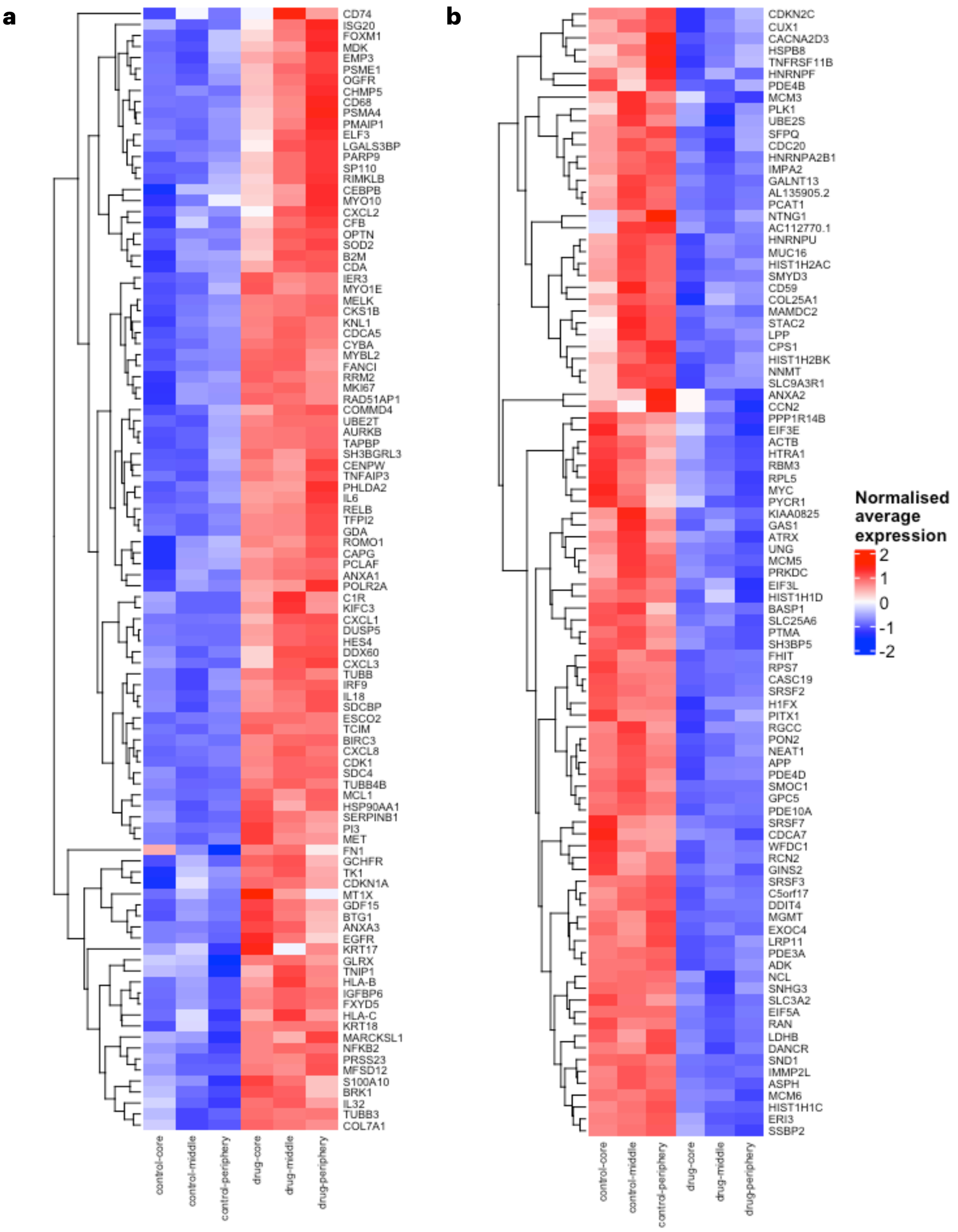

**Supplementary Figure S9.** Normalised average expression levels of differentially expressed genes, filtered for those genes that are affected by irinotecan treatment but do not display a spatially-dependent response. **(a)** Genes that are upregulated on exposure to the drug. **(b)** Genes that are downregulated on exposure.

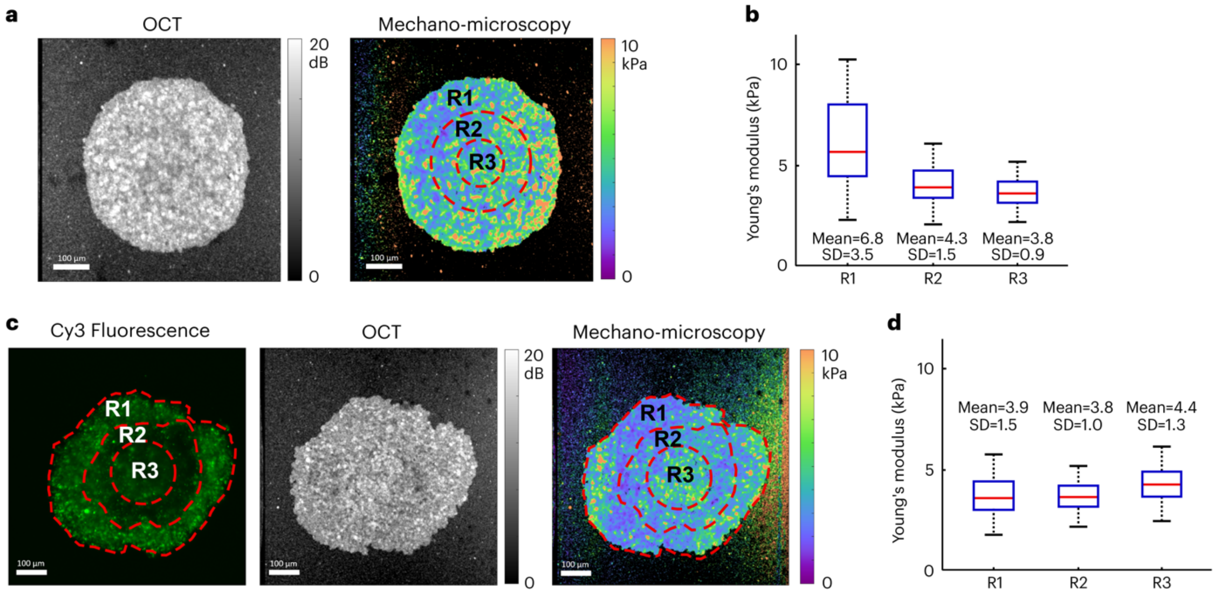

**Supplementary Figure S10.** Layered spheroids display uniform stiffness. **(a)** Optical coherence tomography (OCT) (left) and mechano-microscopy (right) images of mono-layer spheroid. **(b)** Box plot of elasticity measurements for pixels across the three arbitrarily selected regions of interest (R1-3) outlined in (a). Mean and standard deviation (SD) are also shown. **(c)** Fluorescence, OCT and mechano-microscopy images of three-layer spheroid. Cells in the core and outer layers were both tagged with Cy3-SBO to identify boundaries between the different layers since the setup for imaging in conjunction with elastography was a single-colour excitation system. **(d)** Elasticity (Young's modulus) measurements for pixels across the three layers (R1-3) outlined in (c) indicate little change in stiffness between the layers.

### Supplementary Tables

**Supplementary Table S1.** SBO sequences used in this study.

| SBO ID | Feature barcode | Used for labelling |
|---|---|---|
| SBO03 | GTGCAAGAGTTGGCG | 3-layer spheroid, vehicle-core |
| SBO05 | TGGTGACAAGTATCT | 3-layer spheroid, vehicle-middle |
| SBO07 | CGAGGTACATCTTGT | 3-layer spheroid, vehicle-periphery |
| SBO04 | GAAGAAGCGTTATTC | 3-layer spheroid, drug-core |
| SBO06 | CACTCCTTGACAGGT | 3-layer spheroid, drug-middle |
| SBO08 | GCCAAGATCAGGTCC | 3-layer spheroid, drug-periphery |
| SBO01-Cy5 | AAGGCAGACGGTGCA | 2-layer spheroid, periphery |
| SBO14-Cy5 | ACGAATCGGATACTA | 2-layer spheroid, periphery |
| SBO02-Cy3 | GGCTGCGCACCGCCT | 2-layer spheroid, interior |
| SBO13-Cy3 | ATAATCATTACGTGG | 2-layer spheroid, interior |

**Supplementary Table S2.** Raw transcript counts of individual cells labelled either with unmodified SBO or Cy5-conjugated SBO.

| | Min | Q1 | Median | Mean | Q3 | Max |
|---|---|---|---|---|---|---|
| SBO (no fluorophore) | 515 | 16536 | 21494 | 21564 | 26634 | 52496 |
| SBO–Cy5 | 504 | 17338 | 21989 | 23215 | 28808 | 67574 |
| **All cells** | **504** | **17078** | **21788** | **22680** | **27892** | **67574** |

**Supplementary Table S3.** Raw gene counts of individual cells labelled either with unmodified SBO or Cy5-conjugated SBO.

| | Min | Q1 | Median | Mean | Q3 | Max |
|---|---|---|---|---|---|---|
| SBO (no fluorophore) | 169 | 4180 | 4857 | 4597 | 5354 | 6991 |
| SBO–Cy5 | 120 | 4242 | 4849 | 4726 | 5474 | 8489 |
| **All cells** | **120** | **4234** | **4851** | **4684** | **5428** | **8489** |

**Supplementary Table S4.** Raw SBO counts of individual cells labelled either with unmodified SBO or Cy5-conjugated SBO (rounded to the nearest integer).

| | Min | Q1 | Median | Mean | Q3 | Max |
|---|---|---|---|---|---|---|
| SBO (no fluorophore) | 152 | 606 | 772 | 882 | 1030 | 2892 |
| SBO–Cy5 | 256 | 834 | 1370 | 1662 | 2062 | 11450 |
| **All cells** | **152** | **734** | **1090** | **1408** | **1753** | **11450** |

**Supplementary Table S5.** Raw transcript counts of individual cells isolated from two-layer spheroids.

| | Min | Q1 | Median | Mean | Q3 | Max |
|---|---|---|---|---|---|---|
| Interior (Cy5-SBO) | 503 | 12526 | 16148 | 18375 | 22385 | 88622 |
| Periphery (Cy3-SBO) | 507 | 14819 | 19213 | 20901 | 26197 | 58467 |
| **All cells** | **503** | **12951** | **16914** | **19042** | **23434** | **88622** |

**Supplementary Table S6.** Raw gene counts of individual cells isolated from two-layer spheroids.

| | Min | Q1 | Median | Mean | Q3 | Max |
|---|---|---|---|---|---|---|
| Interior (Cy5-SBO) | 339 | 3728 | 4409 | 4452 | 5255 | 7969 |
| Periphery (Cy3-SBO) | 340 | 4192 | 4884 | 4880 | 5721 | 7904 |
| **All cells** | **339** | **3826** | **4530** | **4565** | **5400** | **7969** |

**Supplementary Table S7.** Raw SBO counts of individual cells isolated from two-layer spheroids (rounded to the nearest integer).

| | Min | Q1 | Median | Mean | Q3 | Max |
|---|---|---|---|---|---|---|
| Interior (Cy5-SBO) | 51 | 230 | 564 | 1492 | 1549 | 29972 |
| Periphery (Cy3-SBO) | 67 | 337 | 571 | 1093 | 1149 | 23403 |
| **All cells** | **51** | **262** | **567** | **1386** | **1427** | **29972** |

**Supplementary Table S8.** Differentially expressed genes in two-layer spheroids. Average $\log_2$FC represents expression level in peripheral layer relative to interior.

| Gene | Average $\log_2$FC | *p*-value | $p_{adj}$ |
|---|---:|---:|---:|
| *PGK1* | -0.4164503 | 5.018237e-51 | 1.836725e-46 |
| *NQO1* | 0.3157012 | 4.149931e-48 | 1.518916e-43 |
| *FTL* | 0.2827381 | 3.785542e-47 | 1.385546e-42 |
| *BNIP3* | -0.4473022 | 2.967747e-41 | 1.086225e-36 |
| *AKR1C3* | 0.3987517 | 4.221512e-40 | 1.545116e-35 |
| *ID3* | 0.2783516 | 6.894867e-39 | 2.523590e-34 |
| *CA9* | -0.7937177 | 5.683418e-37 | 2.080188e-32 |
| *FAM162A* | -0.3878044 | 6.854907e-34 | 2.508965e-29 |
| *MYL9* | 0.2556641 | 1.140053e-33 | 4.172708e-29 |
| *GPI* | -0.3431416 | 3.117863e-32 | 1.141169e-27 |
| *LINC01411* | 0.2711669 | 1.517164e-28 | 5.552973e-24 |
| *NDRG1* | -0.2999409 | 5.589421e-27 | 2.045784e-22 |
| *ALDH3A1* | 0.2905408 | 5.520260e-26 | 2.020470e-21 |
| *AKR1C2* | 0.4647274 | 1.071841e-24 | 3.923045e-20 |
| *CCN1* | 0.2505543 | 4.362710e-18 | 1.596795e-13 |
| *KRT17* | 0.2515342 | 4.816368e-14 | 1.762839e-09 |

**Supplementary Table S9.** Raw transcript counts of individual cells isolated from three-layer spheroids.

| | | **Min** | **Q1** | **Median** | **Mean** | **Q3** | **Max** |
|---|---|---|---|---|---|---|---|
| Vehicle | Interior | 2446 | 12552 | 15475 | 17410 | 20456 | 49535 |
| | Middle | 5426 | 11696 | 16695 | 18282 | 21348 | 50713 |
| | Periphery | 1573 | 13590 | 18695 | 21902 | 26674 | 100605 |
| Irinotecan | Interior | 4568 | 14911 | 18802 | 20593 | 24792 | 54274 |
| | Middle | 5215 | 15876 | 19626 | 21743 | 25526 | 69320 |
| | Periphery | 2033 | 17090 | 23168 | 26183 | 32031 | 130156 |
| **All cells** | | **1573** | **15306** | **21011** | **23996** | **29451** | **130156** |

**Supplementary Table S10.** Raw gene counts of individual cells isolated from three-layer spheroids.

| | | **Min** | **Q1** | **Median** | **Mean** | **Q3** | **Max** |
|---|---|---|---|---|---|---|---|
| Vehicle | Interior | 1118 | 3586 | 4235 | 4342 | 5036 | 8603 |
| | Middle | 1973 | 3570 | 4478 | 4527 | 5196 | 7579 |
| | Periphery | 1057 | 3987 | 4820 | 4985 | 5868 | 9176 |
| Irinotecan | Interior | 1881 | 4394 | 5086 | 5084 | 5770 | 8521 |
| | Middle | 2411 | 4570 | 5172 | 5279 | 5909 | 8452 |
| | Periphery | 1211 | 4816 | 5725 | 5716 | 6666 | 9794 |
| **All cells** | | **1057** | **4379** | **5316** | **5370** | **6338** | **9794** |

**Supplementary Table S11.** Raw SBO counts of individual cells isolated from three-layer spheroids (rounded to the nearest integer).

| | | **Min** | **Q1** | **Median** | **Mean** | **Q3** | **Max** |
|---|---|---|---|---|---|---|---|
| Vehicle | Interior | 51 | 93 | 153 | 337 | 295 | 6192 |
| | Middle | 51 | 96 | 140 | 264 | 270 | 3772 |
| | Periphery | 51 | 380 | 1059 | 2483 | 2730 | 46562 |
| Irinotecan | Interior | 53 | 117 | 187 | 304 | 343 | 2625 |
| | Middle | 51 | 126 | 210 | 363 | 363 | 8374 |
| | Periphery | 51 | 481 | 1372 | 2749 | 3518 | 43664 |
| **All cells** | | **51** | **325** | **1009** | **2394** | **2839** | **46562** |
